# GAN-GMHI: Generative Adversarial Network for high discrimination power in microbiome-based disease prediction

**DOI:** 10.1101/2021.07.23.453477

**Authors:** Yuxue Li, Gang Xie, Yuguo Zha, Kang Ning

## Abstract

Gut microbiome-based health index (GMHI) has been applied with success, while the discrimination powers of GMHI varied for different diseases, limiting its utility on a broad-spectrum of diseases. In this work, a generative adversarial network (GAN) model is proposed to improve the discrimination power of GMHI. Built based on the batch corrected data through GAN, GAN-GMHI has largely reduced the batch effects, and profoundly improved the performance for distinguishing healthy individuals and different diseases. GAN-GMHI has provided a solution to unravel the strong association of gut microbiome and diseases, and indicated a more accurate venue toward microbiome-based disease monitoring. The code for GAN-GMHI is available at https://github.com/HUST-NingKang-Lab/GAN-GMHI.

**Importance:** The association of gut microbiome and diseases has been proven for many diseases, while the transformation of such association to a robust and universal disease prediction model has remained illusive, largely due to the batch effects presents in multiple microbiome cohorts. Our analyses have indicated a plausible venue, which is based on GAN technique, towards batch effect removal for microbiome datasets. GAN-GMHI is a novel method built based on the batch corrected data through GAN, as well as GMHI for prediction of a broad-spectrum of diseases.

## Introduction

There are important links between many complex chronic diseases and the human gut microbiome [1]. Specific sets of gut microbes could directly or indirectly influence complex chronic diseases, such as the microbiome dysbiosis in the development of rheumatoid arthritis [2], thus it is natural that the gut microbiome could be utilized for disease prediction [1,3]. However, a general microbiome-based index for the prediction of a broad-spectrum of diseases is lacking.

Previous work has reported the gut microbiome health index (GMHI) [1], a robust index for assessing health status, based on the species-level taxonomic profile of stool metagenomic sequencing samples. GMHI values can be used to classify samples as healthy (GMHI > 0), non-healthy (GMHI < 0), or neither (GMHI = 0), and results in the previous work have shown strong reproducibility on the validation dataset. However, GMHI has limited power to distinguish samples from different diseases: as the stool metagenomes in that study were collected from over 40 published studies, it is nearly impossible to exclude experimental and technical inter-study batch effects [1]. The batch effect refers to the technical difference caused by the processing and measurement of samples in different batches that are not related to any biological variation recorded during the test [4].

A number of traditional tools for batch effect removal have been developed, such as ComBat [5]. However, traditional batch effect removal tools have shown limitations when faced with heterogeneous data of human gut microbiome from hundreds of studies and thousands of individuals. To address this major hurdle, here we introduced GAN-GMHI, based on the generative adversarial network (GAN), for improved discrimination power of GMHI. GAN was applied to reduce the batch effects on a large collection of gut microbiome samples from multiple cohorts containing both healthy and non-healthy individuals. Then GMHI could be applied to the batch corrected data for prediction. Compared with the original GMHI, GAN-GMHI makes the distribution of GMHI values within the group more concentrated and the distinction between healthy and non-healthy samples more clearly. The effectiveness of GAN for cross-cohort batch correction has been demonstrated: the prediction accuracy of GAN-GMHI has been improved to 88.70% for distinguishing healthy individuals and non-healthy individuals, compared to the accuracy of 70.95%, 72.00% achieved by GMHI and ComBat-GMHI. In summary, batch effect does exist in datasets from different sources, and GMHI can better predict the status of health based on GAN corrected datasets.

## Method

The GAN-GMHI framework consists of three stages. First, a dataset containing phenotype and batch information for all samples is constructed. Second, GAN is applied to guide the batch effect correction of raw data. Third, the corrected dataset is output as the training dataset for GMHI prediction (Figure S1).

The batch effect removal method of iMAP [6], a GAN method previously applied on single-cell RNA-Seq data, was adapted for batch effect removal in this study. It is worth noting that the datasets to be batch-corrected by GAN must be classified based on the phenotype first, and the sub datasets of each phenotype are regrouped according to the batch. Such processing ensures that the unwanted technical variations among different datasets are eliminated, and the biological differences between different phenotypes are retained. We should emphasize that GAN-GMHI applied the same core functions as iMAP. However, we optimized the structure of the model to better fit the microbiome abundance data (see the script at https://github.com/HUST-NingKang-Lab/GAN-GMHI/blob/main/scripts/DNN(GAN).ipynb). The model contains a generator and a discriminator. The generator is an autoencoder structure which consists of seven layers and millions of parameters. The discriminator is a dense neural network which consists of three layers. Additionally, we compared GAN with other three batch effect removal tools (ComBat [5], Seurat3 [7], and Harmony [8]) on the same dataset.

## Results

We have performed a comprehensive analysis on the integrated dataset of 2636 healthy and 1711 non-healthy (including 12 disease phenotypes) individuals’ stool metagenomes from 34 published studies [1]. All of these samples are used as a discovery dataset. Additionally, we have used 679 samples (118 healthy and 561 non-healthy) as a validation dataset [1]. The discovery and validation datasets configuration is the same as in Gupta *et al*. [1].

We first assessed and compared prediction results based on the discovery cohort (training data). By comparison of the species-level GMHI before batch effect correction (referred to as RAW), after batch effect correction by GAN, and after batch effect correction by ComBat (**Figure 1**A) for distinguishing samples from healthy and non-healthy individuals, we observed that the prediction accuracy is largely improved after batch effect correction: GAN-GMHI achieved an overall accuracy of 88.70%, which is higher than GMHI’s accuracy of 70.95%. Moreover, batch effect correction by GAN is better than that by other three batch effect correction methods (Tables S1 and S2): GAN-GMHI (accuracy of 88.70%) outperformed ComBat-GMHI (accuracy of 72.00%), Seurat3-GMHI (accuracy of 70.36%), and Harmony-GMHI (accuracy of 44.65%). In summary, we emphasize that GAN reduced batch effect with high fidelity, and augmented the gut microbiome-based health index by profoundly improved discrimination power.

**Figure 1.**
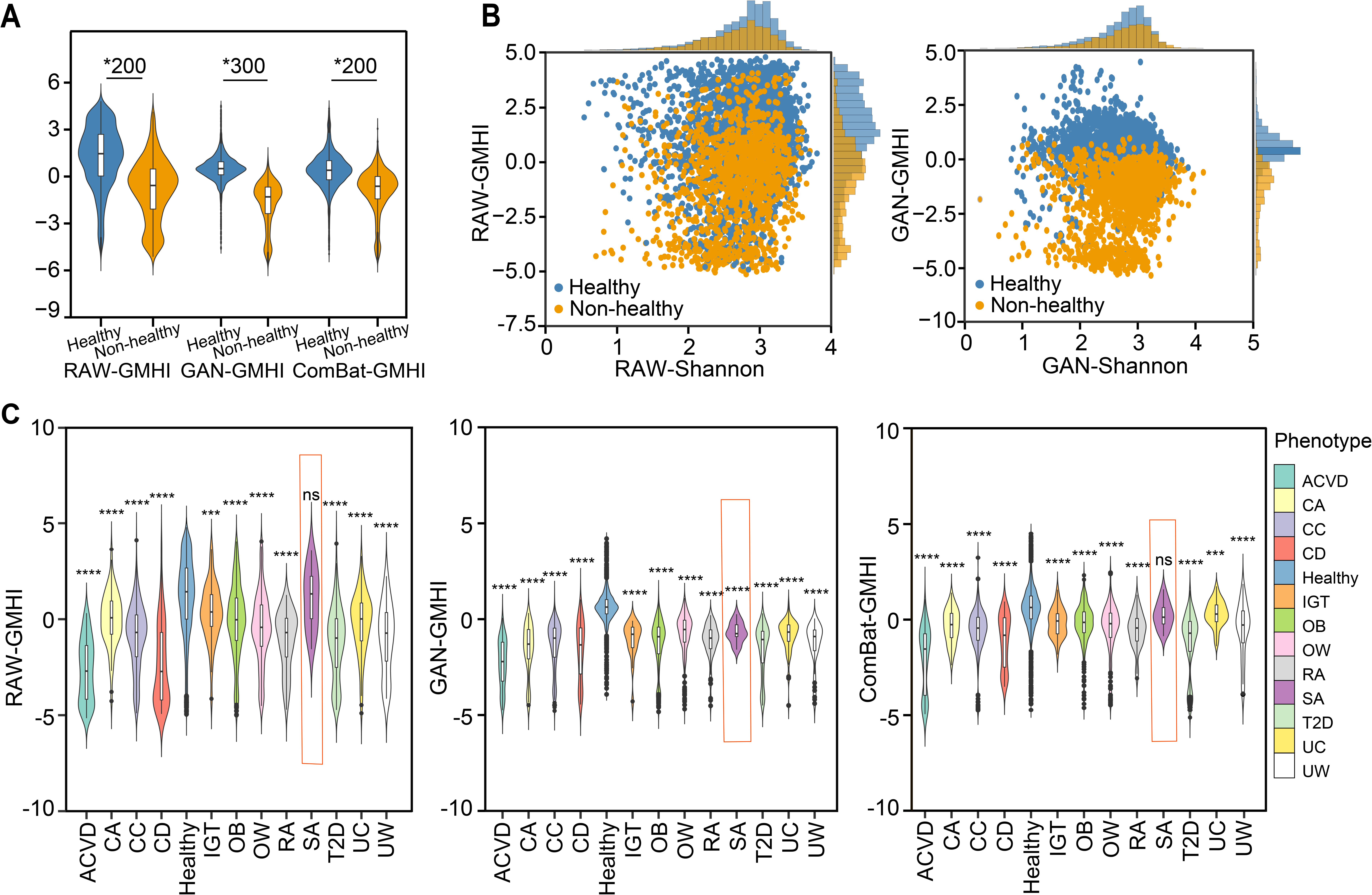
Comparison of GAN-GMHI with other methods under different settings. **A.** Violin plots of GMHI for the healthy and non-healthy groups before (left, referred to as RAW), after batch correction by GAN (middle), and after batch correction by ComBat (right). *200, *P* < 1E-200; *300, *P* < 1E-300; RAW-GMHI, original GMHI; GAN-GMHI, GMHI with GAN enhancement; ComBat-GMHI, GMHI with ComBat enhancement. **B.** the distribution of the RAW (top) and GAN corrected (bottom) GMHI and Shannon diversity. RAW-Shannon, original Shannon index; GAN-Shannon, Shannon index with GAN enhancement. **C.** Violin plots of GMHI index for the healthy and 12 non-healthy phenotypes before (left, referred to as RAW), after batch correction by GAN (middle), and after batch correction by ComBat (right). The red box, an example to show the different results by GMHI, GAN-GMHI, and ComBat-GMHI. ***, *P* < 0.001; ****, *P* < 0.0001; ns, not significant; ACVD, atherosclerotic cardiovascular disease; CA, colorectal adenoma; CC, colorectal cancer; CD, Crohn’s disease; IGT, impaired glucose tolerance; OB, obesity; OW, overweight; RA, rheumatoid arthritis; SA, symptomatic arteriosclerosis; T2D, type 2 diabetes; UC, ulcerative colitis; UW, underweight.

It has been reported that there is a significant change in the alpha diversity (*i.e*., Shannon index) of gut microbiome in non-healthy individuals. Therefore, we compared the abilities of GAN-GMHI and Shannon diversity indicators to differentiate the gut microbiome of healthy and non-healthy individuals. The results demonstrated that GAN-GMHI could yield clearer separation compared with Shannon diversity in differentiating healthy and non-healthy individuals (Figure 1B).

Results on more than ten non-healthy phenotypes have also shown the advantage of GAN-GMHI over GMHI as regard to differentiating these diverse groups of phenotypes. When GMHI was applied, the GMHI values were dispersed over a wide range, and GMHI values for healthy samples were slightly higher than those for non-healthy samples except for symptomatic arteriosclerosis. On the other hand, when GAN-GMHI was applied, the GMHI values were concentrated for each group, and the healthy group was significantly higher than the 12 non-healthy phenotypes (*P* < 0.001 for all non-healthy groups), and the third quartile of GMHI was lower than 0 for all non-healthy phenotypes (Figure 1C). Moreover, it is easier for clinical interpretation based on the results of GMHI. For example, on type 2 diabetes (T2D), GAN-GMHI has captured *Lactobacillus* as biomarkers, which are well founded by published works [9].

Additionally, we compared GAN-GMHI and GMHI on the validation dataset. Cross-cohort batch correction by GAN improved the performance for distinguishing healthy and non-healthy individuals. The prediction accuracy of GAN-GMHI on the validation dataset is 73.05%, compared to the 72.61% of GMHI (Table S1).

Furthermore, GAN is not only applicable for the GMHI disease prediction model, but could also be easily adapted to other models, such as random forest (RF). It has been observed that GMHI and RF exhibit similar performance on the validation dataset, while results of GMHI are easier to interpret clinically (Table S1). We emphasize that although the results of GAN-GMHI and GAN-RF also have similar accuracies on the validation dataset, GAN-GMHI has inherited the interpretability of the GMHI method, and thus is more suitable for clinical interpretation. For example, GAN-GMHI has captured *Lactobacillus* as biomarkers on T2D, which are well founded by published works [9].

Finally, the computational expense of GAN-GMHI is feasible even on a regular laptop. For example, in an experiment including 298 samples from four batches of the colorectal cancer phenotype, the total running time, including training and predicting, is no more than one hour and the maximum usage of memory is less than 2 gigabytes.

## Conclusion

The association of gut microbiome and diseases has been proven for many diseases, while the transformation of such association to a robust and universal disease prediction model has remained illusive, largely due to the batch effects presents in multiple microbiome cohorts. Our analyses have indicated a plausible venue, which is based on GAN technique, towards batch effect removal for microbiome datasets. GAN-GMHI is a novel method built based on the batch corrected data through GAN, as well as GMHI for prediction of a broad-spectrum of diseases. Our study showed that GAN-GMHI could largely reduce the batch effect, and profoundly improved the performance for distinguishing healthy and non-healthy individuals. Batch effect correction by GAN was also better than that by ComBat. In summary, GAN augmented the gut microbiome-based health index, and GAN-GMHI has indicated a more accurate venue towards microbiome-based disease monitoring.

## Code availability

The source code is available at the GitHub (https://github.com/HUST-NingKang-Lab/GAN-GMHI), and BioCode (https://ngdc.cncb.ac.cn/biocode/tools/7275).

## CRediT author statement

**Yuxue Li:** Investigation, Visualization, Writing - Original Draft. **Gang Xie:** Investigation, Writing - Original Draft. **Yuguo Zha:** Writing - Review & Editing. **Kang Ning:** Writing - Review & Editing, Conceptualization, Supervision, Funding acquisition. All authors read and approved the final manuscript.

## Acknowledgments

We are grateful to Mingyue Cheng and Hui Chong for their insightful discussions. This work was partially supported by National Natural Science Foundation of China (Grant Nos. 32071465, 31871334 and 31671374), and the National Key R&D Program (Grant No. 2018YFC0910502).

## Supplementary materials

**Figure S1 Schematic diagram of the technological framework**

**Table S1 The accuracy of the two classifiers for healthy and non-healthy classification**

**Table S2 The overall prediction accuracy of different methods before and after batch correction**

**File S1 Details about the GAN-GMHI method**

