## Supplementary material for "GAN-GMHI: Generative Adversarial Network for high discrimination power in microbiome-based disease prediction": File S1

**File S1 Details about the GAN-GMHI method**

**A data integration approach based on sample phenotypes:**

Transferring the calculation process of a generative adversarial network (GAN) to the data integration of this research, the batch of data with the largest standard deviation needs to be fixed, and other data are regarded as generated data, the difference among fixed data and other batches of data can be minimized at last.

In data integration, the data is first classified by phenotype, and the sub datasets of each phenotype are regrouped according to batches. Batches with less than 15 samples are combined into a reorganization batch, whose sample capacity does not exceed 100, and a recombination batch is formed for excess samples until all samples are merged. It should be noted that the reorganization should be carried out as far as possible in accordance with the principle of "similar batches are merged first", that is, to ensure that the samples in the reorganized batches have larger repetitions or similar properties. Batch differences are mostly based on large sample sizes, and small sample sizes can be ignored, which means that the original data should be kept as much as possible to ensure that the biological significance remains unchanged. Therefore, for some phenotypes, the number of samples in its sub dataset is too small (less than 100 samples) but contains multiple batches, batch correction will not be performed.

For the integration of different batches within the same phenotype, the std function of the python package "NumPy" is used to calculate the standard deviation for descending order. The data with the largest standard deviation is regarded as fixed data, other batches need to be batch corrected by GAN. Choosing the fixed data with the largest standard deviation is beneficial to cover more sample points and types, as far as possible to make the samples of other batches match the fixed sample well, to effectively reduce the batch difference.

**The idea of GAN approach:**

Based on the similarity of data format between the single-cell RNA-sequencing datasets and the microbiome species relative abundance datasets, the batch effect removal method of iMAP [1] was transferred to this study.

**The framework of analysis:**

In this study, a total of metagenomic data from 2,636 healthy and 1,711 non-healthy individuals were collected [2]. Among them, the non-healthy individuals included 12 disease phenotypes. All the data were compiled into meta-datasets as raw data (RAW); The raw data was integrated based on the sample phenotype, and the batch correction data (GAN) was obtained by GAN within the group. Additionally, we also have used 679 samples (118 healthy and 561 non-healthy) as testing set to verify the stability of GAN-GMHI [2].

In order to verify the accuracy and stability of GAN-GMHI, vertical and horizontal comparisons were made in different ways. Firstly, using the RAW and GAN data as the training data sets respectively, and the accuracy of the GAN data set for GMHI and Random Forest prediction has been improved (Table S1). Then, for the two data sets, GMHI and Random Forest methods are used to establish a health prediction model respectively, and make predictions on independent population cohorts (Figure S1). The performance and accuracy of the model trained on the GAN data set is slightly better than that of model based on the unprocessed data (Table S1). On the other hand, we respectively calculated the prediction accuracy of GMHI based on the RAW, GAN, and Combat data sets. The results show that GAN has a more significant improvement in GMHI prediction performance compared to traditional batch correction methods (Table S2).

In general, this study has proved that different batches of intestinal microbiome data have obvious batch effects, and batch correction can reduce the impact of batch effects, providing useful support for health prediction and even biological big data analysis.
