## Supplementary material for "GAN-GMHI: Generative Adversarial Network for high discrimination power in microbiome-based disease prediction": Table S1

**Table S1 The accuracy of the two classifiers for healthy and non-healthy classification**

| **Method** | **Performance on discovery dataset (4347 samples)** | **Performance on validation dataset (679 samples)** |
| --- | --- | --- |
|  | **Overall accuracy (%)** | **Overall accuracy (%)** |
| GMHI | 70.95 | 72.61 |
| RF | 99.71 | 73.17 |
| GAN-GMHI | 88.70 | 73.05 |
| GAN-RF | 99.98 | 75.27 |

*Note*: RF, random forest; GAN-GMHI, GMHI with GAN enhancement; GAN-RF, random forest with GAN enhancement.
