## Supplementary material for "GAN-GMHI: Generative Adversarial Network for high discrimination power in microbiome-based disease prediction": Table S2

**Table S2 The overall prediction accuracy of two methods before and after batch correction**

| **Method** | **Performance on training cohort (4347 samples)** | | | **Performance on testing cohort (679 samples)** | | |
| --- | --- | --- | --- | --- | --- | --- |
|  | **Overall accuracy (%)** | **Accuracy in healthy (%)** | **Accuracy in non-healthy (%)** | **Overall accuracy (%)** | **Accuracy in healthy (%)** | **Accuracy in non-healthy (%)** |
| GMHI | 70.95 | 75.61 | 63.76 | 72.61 | 75.42 | 72.01 |
| GAN-GMHI | 88.70 | 87.03 | 91.29 | 73.05 | 57.63 | 76.29 |
| ComBat-GMHI | 72.00 | 76.10 | 65.63 | 64.06 | 57.63 | 49.02 |
| Seurat3-GMHI | 70.36 | 71.66 | 68.32 | 63.62 | 80.51 | 60.07 |
| Harmony-GMHI | 44.65 | 43.55 | 64.11 | 49.78 | 9.32 | 58.29 |

*Note*: ComBat-GMHI, GMHI with ComBat enhancement; Seurat3-GMHI, GMHI with Seurat3 enhancement; Harmony-GMHI, GMHI with Harmony enhancement.
